# Immunological evaluation in African Green Monkeys of two nasal vaccine candidates to induce a broad immunity against coronaviruses

**DOI:** 10.1101/2025.05.28.656401

**Authors:** L. Lazo, E. Suzarte, D. Vázquez-Blomquist, I. Rodríguez-Alonso, Y. Pérez, P. Puentes, R. Martínez, D. Urquiza, R. Amaya, M. Bequet-Romero, Y. Romero, J. Hernández, R. Chen, W. Li, G. Guillén, Y. Perera, Y. Lobaina, L. Hermida

## Abstract

The COVID-19 pandemic has had a profound impact on the world. While the disease is currently under control, the emergence of new coronavirus variants with pandemic potential underscores the need for more universal coronavirus vaccines. This study evaluates the immunogenicity in non-human primates of two broad-spectrum nasal vaccine candidates called PanCoV. The vaccine preparations are based on highly conserved regions of SARS-CoV-2, specifically the nucleocapsid (N) and the S2 subunit from Spike protein, which include several relevant T-cell epitopes and have the potential to form virus-like particles. PanCoV1 candidate contains the N protein, while the PanCoV2 comprises a chimeric protein containing the C-terminal domain of N protein fused to a fragment of S2 subunit. Both vaccine candidates also include the receptor-binding domain (RBD) from Spike protein and the ODN-39M as CpG adjuvant. The results demonstrate that both nasal vaccine candidates can boost the anti-RBD immune response and induce anti-N immunity, this systemic response shows cross-reactivity with SARS-CoV, MERS-CoV, and H-CoV antigens. In addition, a neutralizing response against SARS-CoV-2 that persisted for almost three months after the last dose was induced. A mucosal-specific and cross-reactive IgA response was also observed, mainly in monkeys vaccinated with PanCoV2. Furthermore, the specific IFNγ-secreting response observed in the vaccinated groups peripheral blood mononuclear cells highlights the ability of these candidates to stimulate cellular immunity with a Th1 bias. PanCoV vaccines emerge as promising candidates for use as a booster alternative, with the potential to amplify and broaden the scope of the immune response against coronaviruses.

## Introduction

The ongoing threat of future pandemics, particularly those caused by coronaviruses, underscores the critical need for proactive vaccine development. Experts projections suggest the occurrence of at least three pandemics as severe as COVID-19 every century, alongside more frequent regional outbreaks [1]. Investing in vaccine research and manufacturing infrastructure before pandemics arise is crucial. This proactive approach involves focusing on high-risk pathogens and creating broad-spectrum vaccines capable of protecting against multiple coronavirus variants. Key strategies under development include mosaic vaccines [2], targeting conserved epitopes [3], leveraging advanced platforms [4,5], enhancing delivery methods [6], and optimizing immune responses through heterologous boosting.

Two nasal-administered protein subunit-based pan-coronavirus (PanCoV) vaccine candidates have recently undergone immunogenicity evaluation in murine models. Both comprising RBD from SARS-CoV-2 Delta strain combined in one case with the nucleocapsid (N) protein from the same variant, while the other includes a chimeric protein known as S2NDH that contains conserved fragments from S2 and N proteins [7,8]. In addition, both candidates include ODN-39M, a CpG oligodeoxynucleotide (ODN), as a Th1-inducing adjuvant [4].

The first candidate based on the nucleocapsid protein, a conserved antigen among coronaviruses shows the formation of aggregated structures, leveraging the natural ability of nucleocapsids to bind nucleic acids [8]. The preparation N + ODN-39M + RBD, administered by intranasal route in BALB/c mice, demonstrated the induction of a robust anti-N cell-mediated immune (CMI) response in the spleen and humoral immunity in both sera and bronchoalveolar lavage fluids (BALF). Notably, the immunity generated was cross-reactive against the N protein from both the SARS-CoV-2 Omicron variant and SARS-CoV. Furthermore, a potent, local and systemic, antibody and CMI response against RBD was generated, demonstrating neutralizing capacity against SARS-CoV-2 and SARS-CoV [8]. On the other hand, the second vaccine candidate includes, as part of a chimeric protein, the C-terminal from N protein and also a fragment from S2 subunit from Spike protein. S2 domain it is recognized by its highly conservation between coronaviruses and has been also pointed out as relevant antigen in the generation of neutralizing antibodies with broader scope of protection [7]. In previous studies was demonstrated that the S2NDH protein preparation form large aggregated structures when mixed with CpG ODN-39M, enhancing its immunogenic potential [2]. Mice immunized intranasally with the bivalent formulation S2NDH + ODN-39M + RBD exhibited significantly elevated levels of anti-RBD IgG and IgA antibodies in both sera and BALF, respectively. Additionally, the formulation elicited strong IgG and IgA responses against the N protein. Furthermore, a positive interferon-gamma (IFNγ) ELISPOT response was recorded against both the conserved peptide N351–365 and RBD. The humoral immune response generated by this bivalent formulation demonstrated cross-reactivity with N proteins from both the SARS-CoV-2 Omicron variant and SARS-CoV. Similarly, anti-RBD antibodies exhibited cross-reactivity with RBD proteins from these variants, indicating a broad protective potential. The neutralizing capacity of this humoral response was also confirmed, supporting this formulation as a vaccine candidate [7].

The aim of the present study is to evaluate the immunogenicity of both bivalent formulations (N + ODN-39M + RBD (PanCoV1) and S2NDH + ODN-39M + RBD (PanCoV2)) in non-human primates, specifically in African Green Monkeys (AGMs). This specie has gained recognition as a valuable model for studying SARS-CoV-2 and assessing COVID-19 vaccines [9–11]. The results obtained show the capacity of these nasal vaccine candidates to boost the anti-RBD immune response and generate anti-N immunity in both mucosal (nasal washes and saliva) and systemic compartments. Cross-reactivity assessments further confirmed the broader antibody responses elicited. In addition, functional analyses indicated that both vaccine formulations induced neutralizing antibody titers against SARS-CoV-2 Delta strain that persisted for several months. Furthermore, the capacity of these nasal candidates to stimulate cellular immunity was also evaluated.

## Materials and methods

### Recombinant proteins and ODN-39M

The recombinant antigens were purchased from Sinobiological Inc. (Beijing, China): Nucleocapsid proteins from SARS-CoV-2 Delta (40588-V07E29), from SARS-CoV (40143-V08B), MERS-CoV (40068-V08B) and HCoV-229E (40640-V07E), S2 protein from SARS-CoV-2 Ancestral strain (40590-V08H1), from SARS-CoV (40150-V08B3) and MERS-CoV (40070-V08B) and RBD protein from SARS-CoV-2 Delta (40592-V08H90) and SARS-CoV (40150-V08B2).

S2NDH, a chimeric recombinant protein with a molecular weight of 40 kDa, was produced in *Escherichia coli*. It was purified using SP Sepharose fast-flow and immobilized metal affinity chromatography (IMAC) techniques in the presence of the chaotropic agent urea at a concentration of 8 M. The renatured protein achieved an estimated purity level exceeding 95% [7].

The ODN-39M, a 39 mer, whole-phosphodiester backbone oligodeoxynucleotide (5′ -ATC GAC TCT CGA GCG TTC TCG GGG GAC GAT CGT CGG GGG-3′), was synthesized by Sangon Biotech (China)[12].

### Vaccine candidate preparations

The chimeric protein S2NDH was incubated with the ODN-39M. Briefly, in a 100 µL reaction, 50 µg of S2NDH were mixed with 60 µg of ODN-39M in buffer 10 mM Tris, 6 mM EDTA, pH 6.9. Six independent reactions were prepared and incubated for 30 min at room temperature (RT) (20–25°C) and then stored at 4°C for 4h. The final dose per animal contained 62.5 µg of the recombinant protein and 75 µg of ODN-39M.

The N protein from SARS-CoV-2 Delta variant, was subjected to *in vitro* aggregation, as previously described [12,13], with few modifications. Briefly, in an 800 µL reaction, 320 µg of N protein was mixed with 480 µg of ODN-39M in Tris/EDTA buffer, pH 6.9. The mixtures were incubated for 15 min at RT. Finally, the preparation was centrifuged at 14,000 × g for 10 min. The resulting supernatant was collected and tested for protein concentration. The final dose per animal contained 50 µg of the recombinant protein and 75 µg of ODN-39M.

Placebo formulation contained 75 µg of ODN-39M and buffer. Independent formulations were prepared for each immunization point.

### Animals and ethics statement

Healthy male adult (3-7 years old and weight of 5.5 ± 0.5 kg) African green monkeys (*Cercopithecus aethiops sabaeus*) were obtained from the Center for the Production of Laboratory Animals (CENPALAB, Cuba). The study was carried out in strict accordance with the recommendations of the Guide for the Care and Use of Laboratory Animals of the Center for Genetic Engineering and Biotechnology (CIGB, Cuba). The animals were maintained in individual cages throughout the study and were fed with commercial monkey chow supplemented with fruits and vegetables. Water was available *ad libitum*. The monkeys were subjected to clinical inspections of lymph nodes, skin, respiratory, digestive, and nervous systems. They did not show any signs of pain or distress at any time during the study. All results were reported and records were maintained in accordance with CIGB and Cuban guidelines for animals used in research. Temperature and body weight of all animals were assessed by veterinary staff. Clinical biochemistry assays were performed monthly to minimize animal stress. All immunizations and fluid samples collections were performed using sedation by intramuscular administration of ketamine hydrochloride, 10 mg/kg body weight, and all efforts were made to minimize suffering. To provide variety in their environment, toys and branches were placed inside the cages. None of the monkeys was sacrificed. Before the beginning of the study, all animals were screened by ELISA for prior exposure to SARS-CoV-2.

### Immunization study design and sample collections

Nine male African green monkeys were randomly divided into three groups of three animals each (Figure 1a). They were immunized intranasally with three preparations as follows: group 1, PanCoV1 (containing 50 µg of N, 75 µg of ODN-39M and 50 µg of RBD); group 2, PanCoV2 (containing 62.5 µg of S2NDH, 75 µg of ODN-39M and 50 µg of RBD); and group 3, placebo (containing 75 µg of ODN-39M). The doses were administered, with a 200uL pipette, at days 0, 30, and 90, using a total volume of 200 µL/dose/animal (100 µL per nostril).

**Figure 1.**
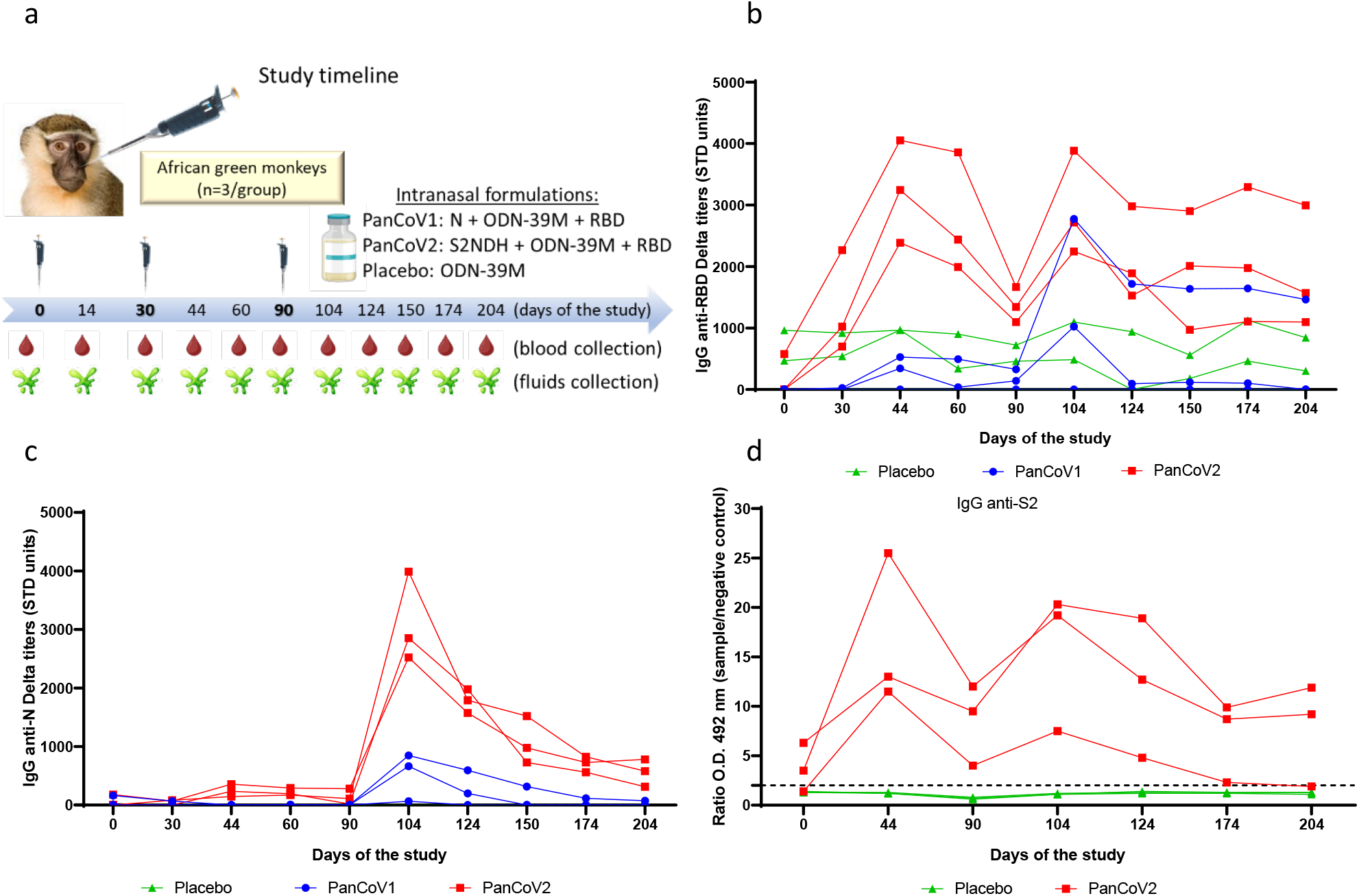
Kinetic of IgG immune response elicited in plasma after vaccination. African green monkeys were intranasally immunized with three doses (0, 30, and 90 days) of each vaccine candidate or Placebo. General representation of the study, chronogram of doses and samples collection (a). IgG antibody titers were measured in plasma by ELISA: (b) against RBD from Delta variant, (c) against N protein from Delta variant, and (d) against S2 protein from Ancestral variant. STD refers to standard. The dotted line indicates the positive cutoff (ratio OD 492 nm sample/background control on the same plate equal to 2).

Fluid samples were collected at days 0, 14, 30, 44, 60, 90, 104, 124, 150, 174 and 204 and were stored at −80°C. Nasal washes were obtained instilling 500 µL of phosphate-buffered saline (PBS) per nostril and allowing the fluid to drain into a petri dish. Saliva was collected, under the effects of ketamine, by gravity falling into a petri dish without using any salivating agent.

A total of 8-10 mL of blood was collected from the femoral vein using a sterile syringe. Blood samples were immediately transferred into VACUETTE® tubes containing K3EDTA as an anticoagulant. PBMCs were isolated from whole blood using Ficoll density gradient centrifugation. Blood samples were first diluted with an equal volume of PBS. The diluted blood was carefully layered over Ficoll-Paque™ in a 50 mL conical tube, ensuring minimal mixing at the interface. The tubes were centrifuged at 400 × g for 30 minutes at RT without brakes to allow the formation of distinct layers. Following centrifugation, the 1:2 diluted plasma was collected and aliquoted into clean polypropylene tubes, and stored at −80°C for further analysis. The PBMC layer, located at the plasma-Ficoll interface, was carefully aspirated using a sterile pipette and transferred to a new tube. The collected cells were washed twice with cold RPMI-1640 medium by centrifuging at 300 × g for 10 minutes. Following isolation, a portion of the PBMCs was used fresh in ELISpot experiments, while the remaining cells were frozen to perform the interferon-gamma secretion kinetics. PBMCs were resuspended in freezing medium consisting of 90% fetal bovine serum (FBS) and 10% dimethyl sulfoxide (DMSO) at a concentration of 1–10 × 10^6^ cells/mL. Aliquots of 1 mL were transferred into cryovials, which were placed in a controlled-rate freezing container (Mr. Frosty, Thermo Fisher Scientific) and stored at –80°C overnight. After 24 hours, cryovials were transferred to liquid nitrogen storage for long-term preservation.

### Humoral immune responses

The systemic and mucosal antibody response was evaluated by ELISA. Anti -IgG and -IgA ELISAs were conducted as previously described [7]. Briefly, recombinant protein was used to coat 96-well high-binding plates (Costar, USA) (5 µg/mL for N Delta, 2 µg/mL for RBD Delta, RBD SARS-CoV, N SARS-CoV, N MERS and N HCoV-229E, and 3 µg/mL for S2 ancestral, S2 SARS-CoV and S2 MERS). Plates were subsequently blocked with 2% BSA in PBS with 0.05% Tween 20. Samples were added in duplicates, starting from 1:200 dilution in the case of plasma for homologous antigens (1:50 for heterologous antigens and S2 ancestral) whereas saliva and nasal washes were assayed 3-5 times diluted or non-diluted, respectively. Specific horseradish peroxidase conjugates (Sigma, USA) were used. As a substrate, OPD (Sigma, USA)/hydrogen peroxide solution was employed. Plates were incubated for 10 min in the dark, and the reaction was stopped by the addition of 12.5% H_2_SO_4_. The optical density (OD) was read at 492 nm in a multiplate reader (SUMA, Cuba). The arbitrary units of total IgG titers (anti-RBD and anti-N of SARS-CoV-2, anti-RBD and anti-N of SARS-CoV and anti-N of MERS-CoV and H-CoV) were calculated by plotting the OD values obtained for each sample in a standard curve (hyper-immune serum of known titer). The positivity cutoff was established as twice the average of standard units obtained for the placebo group. For assessments where a standard curve was not available, the results were expressed as the ratio between the sample’s OD and the OD obtained for the background control on the plate (dilution buffer). In this case, for IgG anti-S2 from MERS-CoV and SARS-CoV, and IgA anti-RBD and anti-N from SARS-CoV, the positivity cut-off was established as twice the average value obtained for the placebo group. On the other hand, for kinetic profile evaluations (IgG anti-S2 of SARS-CoV-2 and IgA anti-RBD and N of SARS-CoV-2), the cut-off was defined as a ratio equal to 2.

### Viral neutralization

The assay was carried out in a reference BSL-3 laboratory (Civil Defense lab, Cuba) as has been previously reported [14]. Briefly, 96-well culture plates seeded with 2×10^4^ cells/ well of VERO E6 cell line expressing the ACE2 receptor were used. Cells were incubated for 24 h at 37°C and 5% CO_2_ to form a monolayer grown to 85-90% confluency. Subsequently, the previously inactivated (by incubation at 56°C for 30 minutes) plasma was serially diluted 1:2 in a minimal essential medium (MEM, Gibco, UK) with 2% (v/v) fetal bovine serum (FBS, Capricorn, Germany). Each dilution was incubated for 1 h at 37°C with 100 TCID50 (50% of Tissue Culture Infectious Dose) of SARS-CoV-2 (hCoV-19/CUBA/DC05/2021 (Delta) strain). Viral suspensions in the presence and absence of experimental plasma were used to infect cell monolayers for 96 h at 37°C and 5% CO_2_. After this time, the presence of the cytopathic effect (viral plaques) was verified with the help of an optical microscope and staining with neutral red. The excess vital dye was removed by three washes, and the neutral red incorporated into the cells was dissolved by incubating for 15 minutes at 25°C with a lysis solution composed of 50% ethanol and 1% acetic acid. The absorbance was measured at 540 nm, and the neutralizing titer was reported as the highest dilution of the plasma with an absorbance greater than the cut-off value of the assay. The cut-off value was defined as half of the average absorbance values of the wells without virus.

### Cell mediated immunity

The ELISPOT assay was conducted using monkey IFNγ ELISPOT PLUS (HRP) kit (Mabtech). The 96 well polyvinylidene plates were pre-coated with anti-IFNγ monoclonal antibody (mAb MT126L). After washing with PBS and blocking (medium with 10% FBS) the plates, 200 000 fresh (or thawed) PBMCs per well were plated in duplicate. Cells were stimulated by using different coronavirus viral proteins (10 ug/mL) and incubated for 96h at 37°C. Subsequently, plates were washed with PBS before the addition of biotinylated anti-IFNγ MAb (7-B6-1) at 1 μg/mL. Following incubation at RT for 2h, plates were washed with PBS and incubated with the manufacturer’s recommended dilution of streptavidin-conjugated HRP for 1h at RT. Following washes with PBS, spots resulting from individual IFNγ-producing cells were visualized after a 15-min reaction with TMB ready to use substrate solution, counted with an ELISpot reader (Mabtech IRIS 2). As negative controls, cells were plated without stimulus, and as positive control cells were incubated with ConA (Sigma, USA) at final concentration of 5ug/mL. Positive response was expressed as the average spot count in antigen stimulated wells after subtracting average spots resulting from non-stimulated cells.

### Statistical analysis

GraphPad Prism software v8.0.2 (GraphPad Software, Boston, MA) was used for most of the figure representations. Inferential statistical analyses were not performed due to the small number of animals per treatment group used in the study.

## Results

### Systemic humoral immune response

The specific humoral immune response induced in plasma, was assessed by ELISA. Figure 1b illustrates the kinetic of IgG antibody response anti-RBD Delta from day 0 to day 204 of the study. On the evaluation corresponding with day 0 we found some animals (3 out 9) with positive levels of anti-RBD IgG antibodies. Fourteen days after the second and third doses, a booster response was clearly observed for most animals receiving vaccine candidates (greater than a fourfold increase in titers compared to those detected on day 0 of the study). One animal from PanCoV1 group remains without seroconversion during all the study. The level of the booster response shows a trend to be higher in the animals immunized with PanCoV2. In the majority of vaccinated animals, anti-RBD antibody titers remained positive for nearly four months after the third dose, with titers exceeding 2.8 times those detected in the placebo group (3 out of 3 animals in the PanCoV2 group and 1 out of 3 in the PanCoV1 group).

Unlike the IgG anti-RBD response, no antibody titers against the N protein were detected at day 0 of the study (Figure 1c). The highest response was observed fourteen days after the third dose in 5 out of 6 vaccinated monkeys, with a trend to develop higher titers in the animals receiving PanCoV2 candidate. In 4 out of 6 vaccinated animals the IgG anti-N response remains positive until the last evaluated timepoint on day 204 of the study (IgG anti-N titers ranged from 74.8 to 781.7 in vaccinated animals, compared to 0 in the placebo group) Additionally, the IgG response anti-S2 was evaluated for the PanCoV2 group in comparison to Placebo group (Figure 1d). On day 0 some animals (2 out 6) showed a low positive response. In all PanCoV2 vaccinated monkeys the anti-S2 IgG response was reinforced after each dose, resulting in a positive response detected in all animals up to day 174. In this case, 2 out of 3 animals receiving PanCoV2 show a long-lasting IgG response.

Furthermore, the potential cross-reactivity, against betacoronavirus and alphacoronavirus antigens, of the humoral immune response elicited after intranasal administration of the vaccine candidates was also assessed (Figure 2).

**Figure 2.**
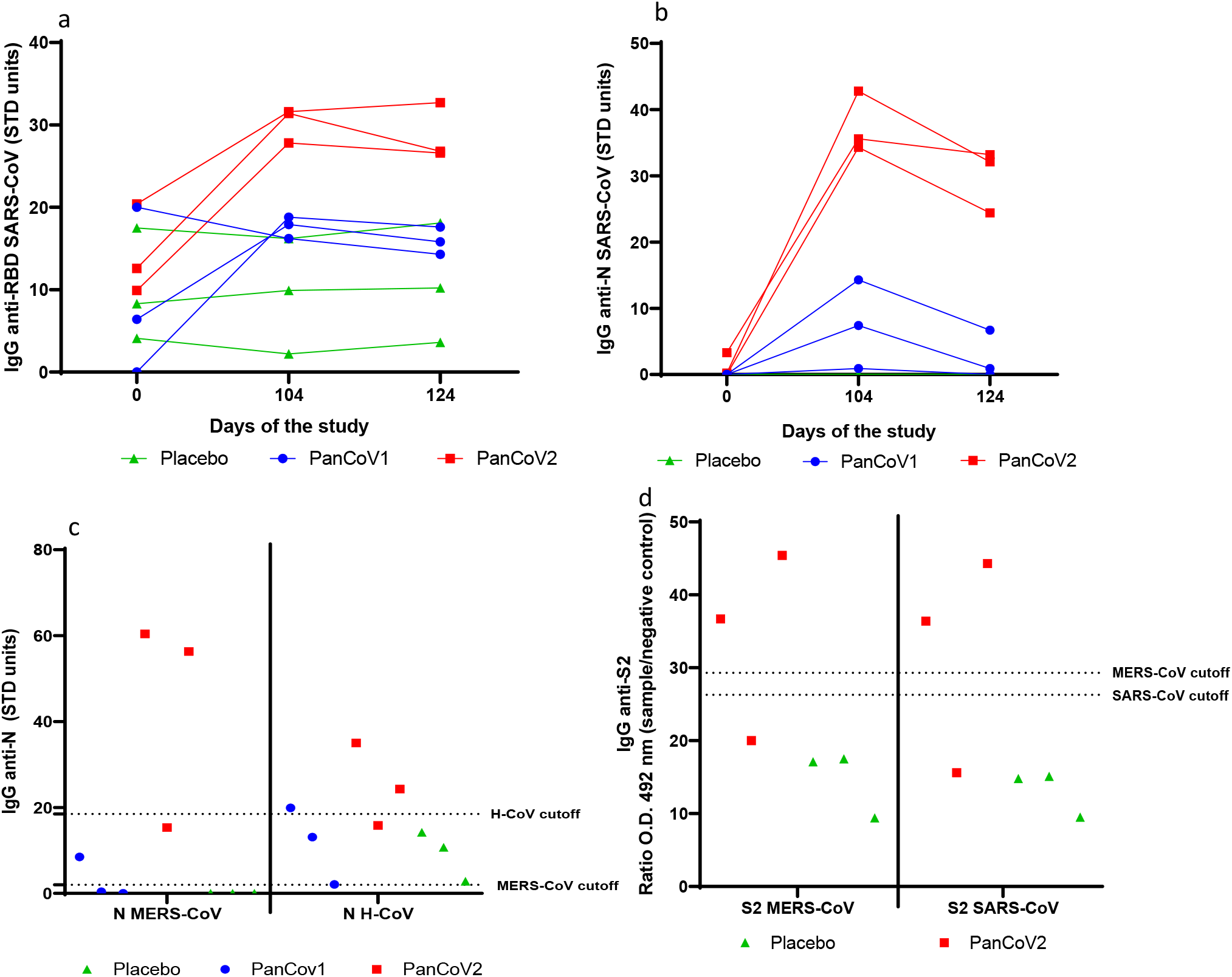
Cross-reactive IgG immune response elicited in plasma after vaccination. African green monkeys were intranasally immunized with three doses (0, 30, and 90 days) of each vaccine candidate or Placebo. IgG antibody titers were measured in plasma by ELISA: (a) against the RBD of SARS-CoV (b) against N protein from SARS-CoV, at 0, 104 and 124 days; and at day 104 (c) against N protein from MERS-CoV and H-CoV-229E, (d) against S2 of SARS-CoV and MERS. STD refers to standard. The dotted line indicates the positive cutoff (twice the average data of the placebo group).

RBD and N proteins from the SARS-CoV, N protein from MERS-CoV and H-CoV-229E and S2 protein from SARS-CoV and MERS-CoV were used as coating antigens in the ELISAs. At day 0 of the study, 8 out of 9 animals showed a positive response against the RBD from SARS-CoV, while only one exhibited a response against the N proteins at that time point. Five out six vaccinated animals show a reinforce of the crossreactive IgG response against the SARS-CoV antigens on day 104 (Figure 2a, b).

A trend to generate a higher response against both SARS-CoV antigens was observed in the animals immunized with PanCoV2 candidate, persisting even one month after the last dose (Figure 2a, b). Similarly, the IgG responses against the N protein from MERS-CoV and H-CoV-229E tend to be higher in the PanCoV2 group, where two out of three animals tested positive, compared to only one animal in the PanCoV1 group (Figure 2c). In addition, positive IgG responses anti-S2 from SARS-CoV and MERS were observed in two animals of PanCoV2 group (Figure 2d).

### Antibody response in naso-pharyngeal mucosa and saliva

The evaluation of the mucosal humoral response was conducted using nasal washes and saliva (Figure 3). Fourteen days after the second dose, a positive IgA anti-RBD response was detected in the nasal washes of four out of six monkeys immunized with the vaccine preparations (two receiving PanCoV2 and two of PanCoV1 group) (Figure 3a). After the third dose of vaccine (administered on day 90) a marked increase in the IgA response was observed in 3 out 6 vaccinated animals and these antibodies remain at high levels until last evaluation on day 204. Additionally, on day 124, four vaccinated animals showed also a positive IgA anti-RBD response in saliva (Figure 3c).

**Figure 3.**
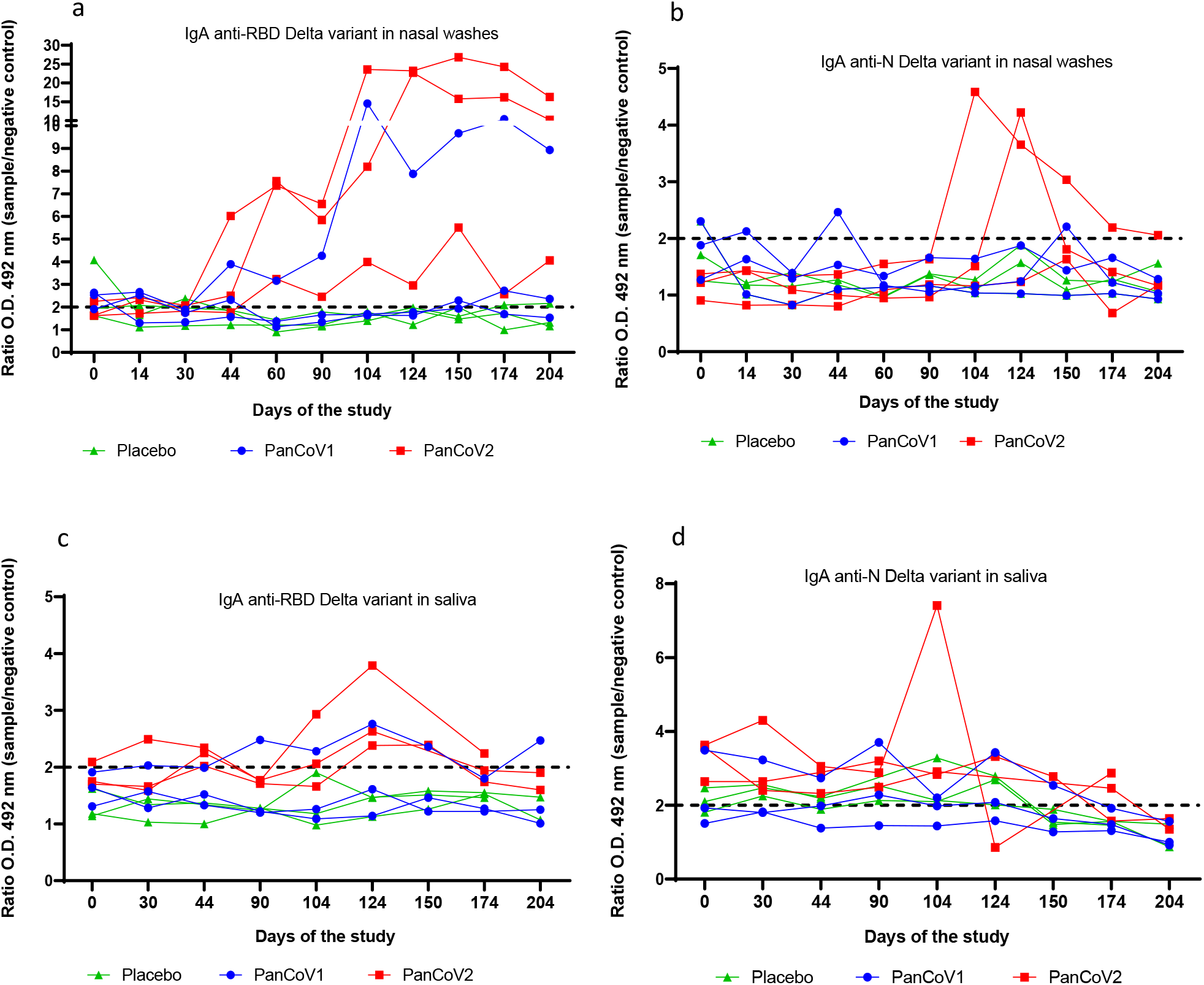
Kinetic of IgA response in nasal washes and saliva. African green monkeys were intranasally immunized with three doses (0, 30, and 90 days) of each vaccine candidate and Placebo. IgA antibody levels were measured by ELISA: against the RBD from Delta variant in non-diluted nasal washes (a) and 1:5 diluted saliva (c); against N protein from Delta variant in non-diluted nasal washes (b) and 1:3 diluted saliva (d). The dotted line indicates the positive cutoff (ratio OD 492 nm sample/background control on the same plate equal to 2).

Although the observed IgA anti-N response in nasal washes tended to be lower compared to the anti-RBD response, two animals in the PanCoV2 group exhibited a positive response converging on day 124; one of these animals maintained that positivity throughout the experiment (Figure 3b). In saliva, only one animal from PanCoV2 group shows a clear positive anti-N response on day 104 (Figure 3d).

The IgA response specific against S2 protein was measured in nasal washes from PanCoV2 immunized monkeys, but no positive response was found (data no shown).

The cross-reactive capacity of the IgA antibody response generated in nasal washes against the RBD and N proteins from SARS-CoV was assessed by ELISA 14 days after the third dose (Figure 4). Two monkeys in the PanCoV2 group, and one in the PanCoV1 group, out of three exhibited a positive recognition of the RBD from SARS-CoV, while the recognition of N protein was detected only in the two animals from PanCoV2 vaccinated group.

**Figure 4.**
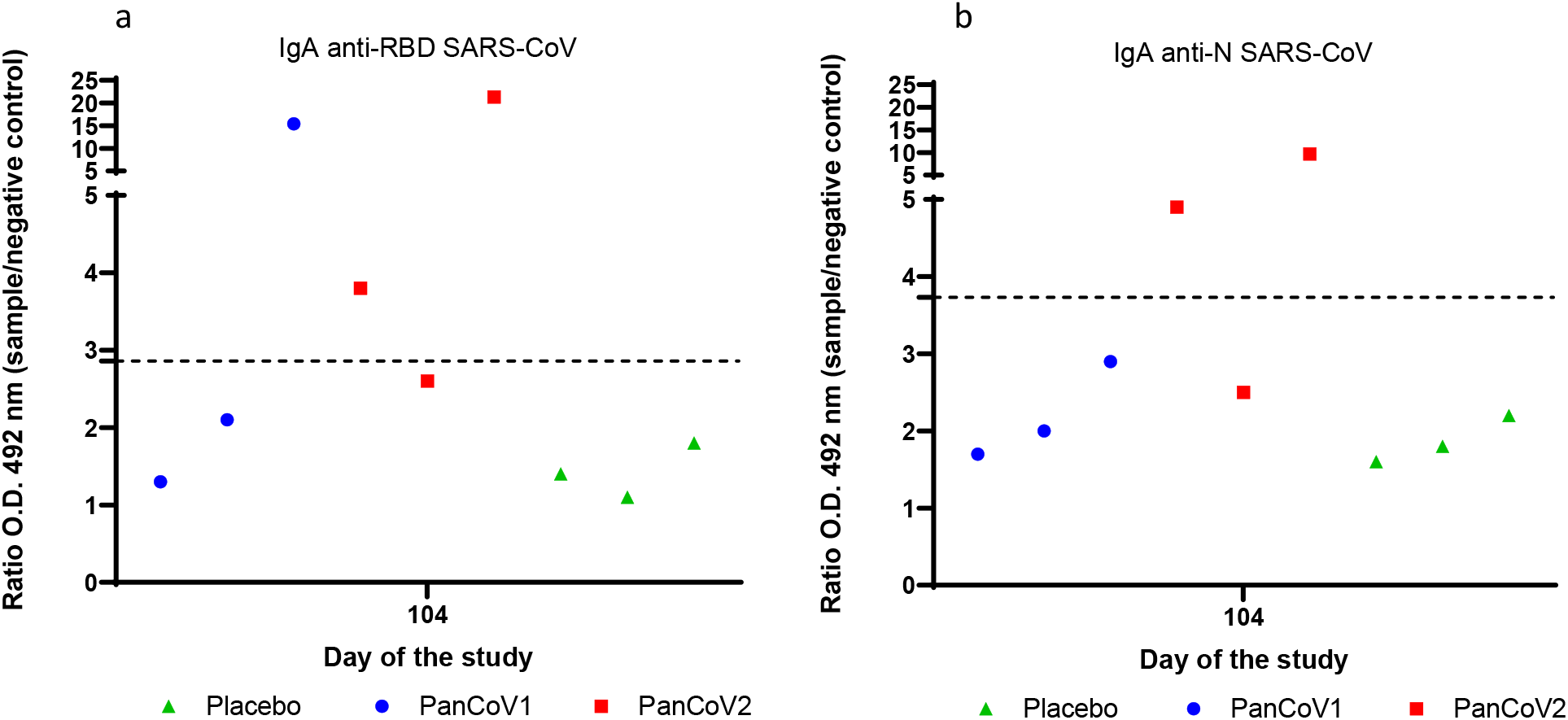
Cross-reactive IgA response elicited in nasal washes. African green monkeys were intranasally immunized with three doses (0, 30, and 90 days) of each vaccine candidate and Placebo. IgA antibody levels were measured by ELISA in nasal washes (non-diluted samples) collected at day 104 (14 days after third dose): (a) against the RBD from SARS-CoV, (b) against N protein from SARS-CoV. The dotted line indicates the positive cutoff (twice the average data of the placebo group).

### Neutralizing antibody response

To assess the functionality of the generated anti-RBD IgG antibody response, plasma from individual monkeys were evaluated using a classical neutralization assay against the SARS-CoV-2 Delta variant (Figure 5). All animals receiving the nasal vaccine candidates in study showed neutralizing antibody titers on day 104, which persisted through day 174, nearly three months after the last dose. At both time-points, the animals immunized with PanCoV2 candidate show a trend to elicit higher neutralizing titers. This contrasts with only one animal in the placebo group exhibiting a positive neutralizing antibody titer (1:56) on day 174.

**Figure 5.**
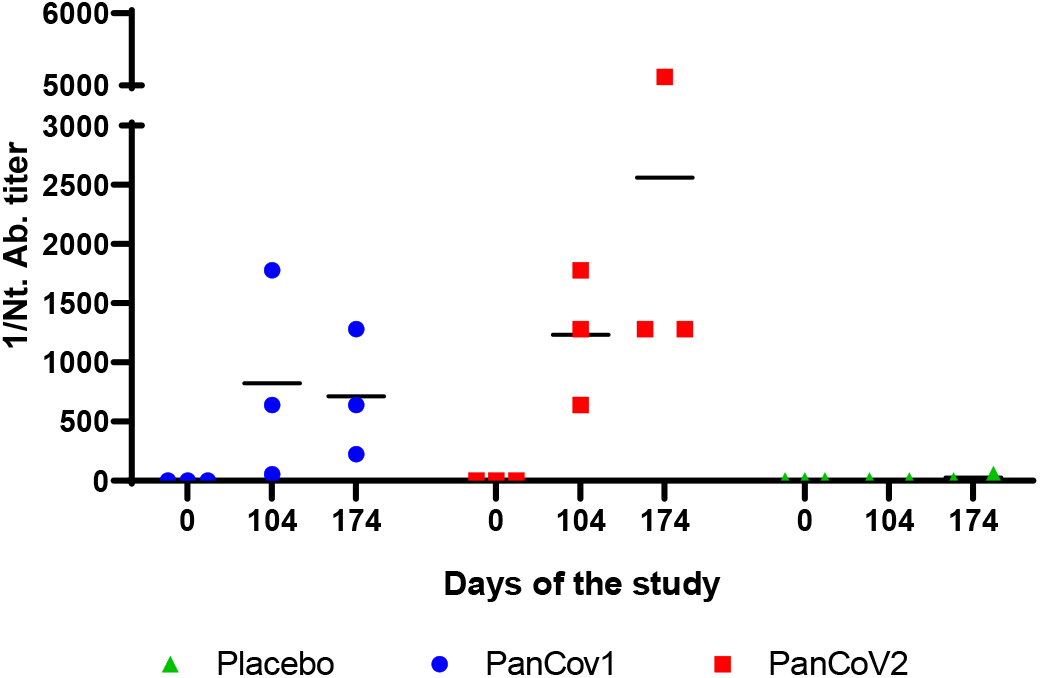
Neutralizing antibody response against SARS-CoV-2 Delta variant measured in plasma. African green monkeys were immunized with three doses (0, 30, and 90 days) of each vaccine candidate or Placebo by intranasal route. Neutralizing antibody titers were measured, in individual plasma samples collected at different time points (0, 104 and 174 days), using a microneutralization assay.

### Cellular immune response

To evaluate the cellular immune response induced by the vaccine preparations, PBMCs from all animals were purified 34 days after the last dose (day 124). These fresh isolated PBMCs were then *in vitro* stimulated with the three proteins included in the vaccine candidates (Delta variant). The number of IFNγ-secreting cells per million of PBMC was determined using an ELISPOT assay (Figure 6).

**Figure 6.**
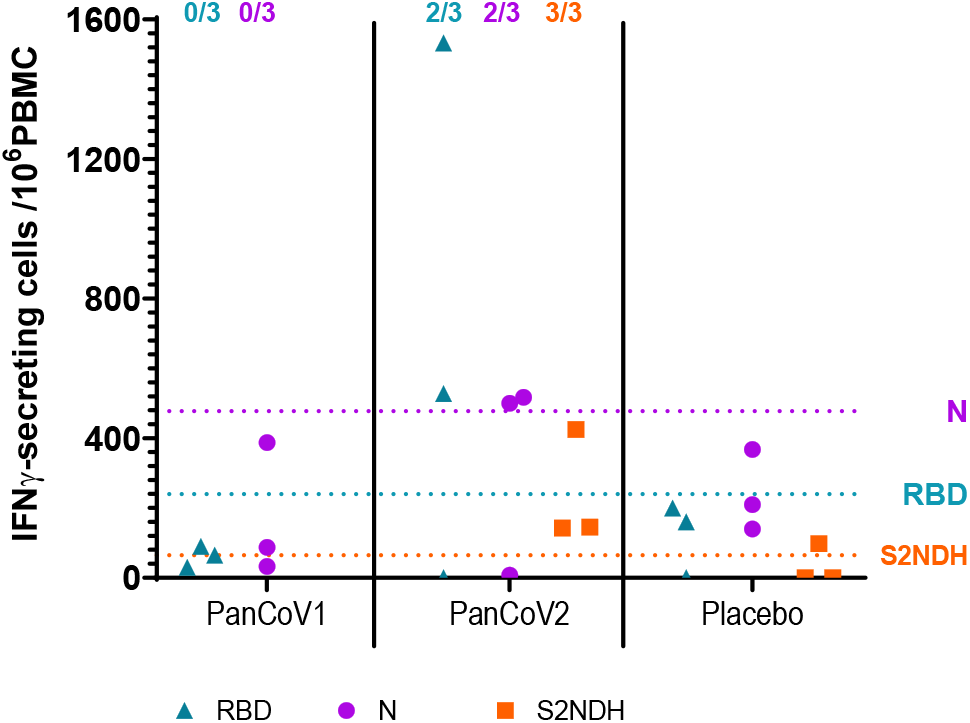
IFNƴ-secretion response measured by ELISPOT in fresh-isolated PBMC. African green monkeys were intranasally immunized with three doses (0, 30, and 90 days) of each vaccine candidate or Placebo. At day 124 of the study, corresponding with 34 days after the third dose, PBMCs were purified and cultured in the presence of protein stimuli: N and RBD from SARS-CoV-2 Delta strain, or S2NDH chimeric protein. The dotted line represents the cutoff value, defined as two times the average response observed in the placebo group following stimulation with each respective protein. The numbers above the graph refer to positive responders in relation to the total number of animals per treatment group.

As illustrated in Figure 6, the frequency of IFNγ-secreting cells was higher in the PanCoV2 formulation when PBMCs were stimulated with RBD and N proteins compared to the PanCoV1 group. Although the response to stimulation with the chimeric protein S2NDH was lower, all animals in the PanCoV2 group still exhibited a positive response.

Another IFNγ ELISPOT evaluation performed using thawed PBMC, corroborated the generation of a cellular immune response against RBD protein (Figure 7). Two out of three animals in the PanCoV1 group showed an increase in the number of IFNγ-secreting cells on day 124 compared to day 0, which was higher than the double of the maximum increase achieved in the placebo group. On the other hand, two out of three animals immunized with PanCoV2 showed an increase in the response on day 124 that persisted until approximately three months after the last administration of the nasal vaccine.

**Figure 7.**
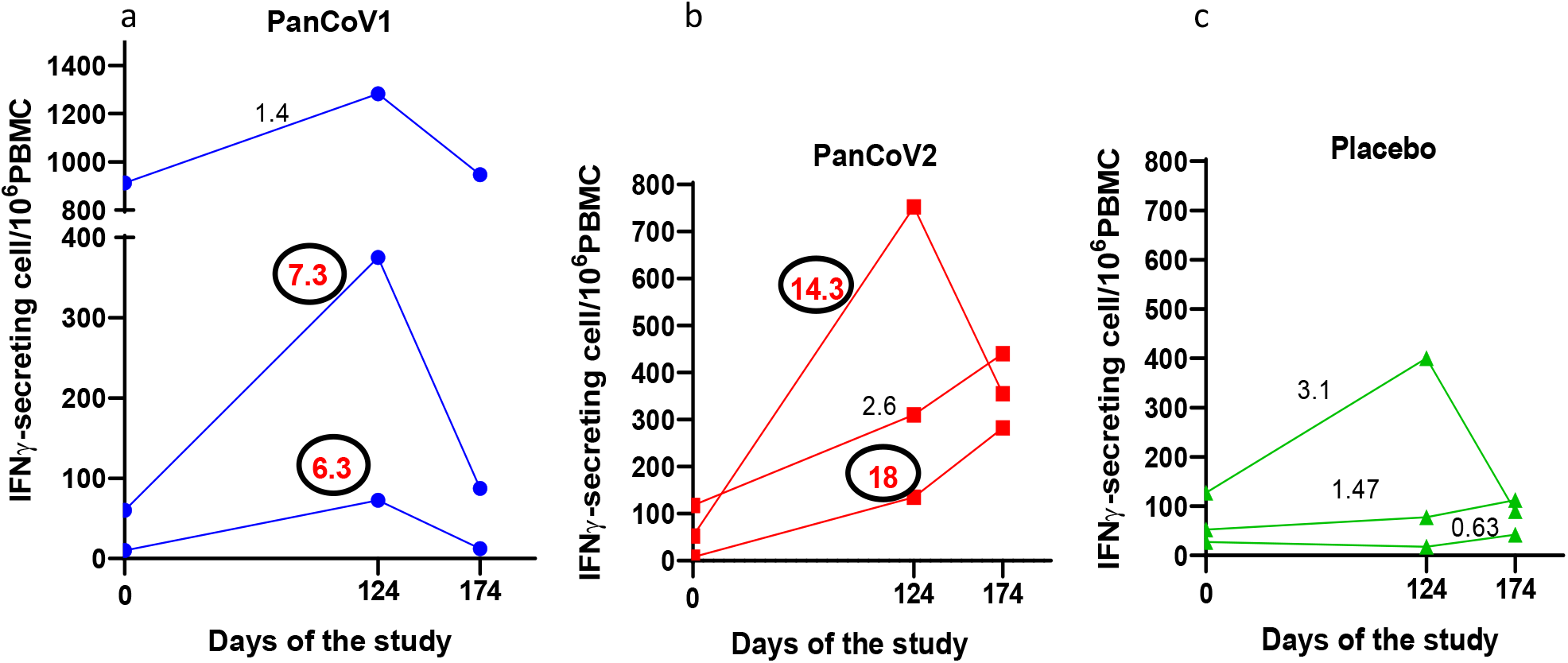
Kinetic of IFNƴ-secretion response measured by ELISPOT in thawed PBMCs. African green monkeys were intranasally immunized with three doses (0, 30, and 90 days) of PanCoV1 (a), PanCoV2 (b) and placebo (c) preparations. PBMCs purified and frozen on days 0, 124 and 174, were thawed and cultured in the presence of RBD protein from SARS-CoV-2 Delta strain. The numbers above the graphs represent the fold increase in the number of IFNγ-secreting cells on day 124 relative to time 0 for each animal. Circled numbers highlighted in bold red denote animals considered positive (double the maximum increase detected in the placebo group).

## Discussion

Considering the possibility of emergence of new coronavirus variants with pandemic potential, the development of more universal vaccines is essential to be prepared against future pandemics threats and ensure a resilient healthcare system. Next-generation vaccines must overcome the limitations of current vaccines by inducing mucosal immunity and long-lasting and broader protection against new strains.

Currently, UB-612 constitutes one of the most advanced next-generation COVID-19 vaccines. This protein/peptide subunit vaccine is in Phase III trials, and was designed as a heterologous booster targeting multiple SARS-CoV-2 proteins with the aim of elicit robust B-cell and T-cell immunity against prototype SARS-CoV-2 viral strains and multiple virus of concern [10]. Another candidate in Phase III trial is CoviLiv™, a live-attenuated intranasal vaccine, which use synthetic biology to re-code SARS-CoV-2’s genetic material [11]. GEO CM04S1 is other synthetic attenuated vaccine, but using a modified vaccinia virus Ankara vector to express spike and nucleocapsid proteins, aiming to generate humoral and cellular immune responses; it is currently in Phase II trials in the U.S. Similarly, VXA-CoV2-1 and VXA-CoV2-1.1-S are composed of the genes coding for both the SARS-CoV-2 S protein and the N protein. These vaccines candidates, currently in phase I and II clinical trials, are formulated as oral tablets enteric coated for efficient delivery to the small bowel and for inducing an IgA response in respiratory tracts [12]. These and other novel strategies being developed include antigenic regions different from the spike protein or multiple RBDs. Furthermore, alternative administration routes are being used to generate immune memory responses localized in the respiratory tract and prevent infection, dissemination, and transmission of the virus.

In this study the immunogenicity of two broad-scope nasal vaccine candidates, called PanCoV, has been evaluated in non-human primates. The previous development and evaluation in mice of these two candidates have been recently published by our group [7,8]. Briefly, the strategy followed for the design of both vaccine preparations was based on the selection of highly conserved regions from SARS-CoV-2 virus, including several relevant T cell epitopes and with the potential to associate with CpG-ODN adjuvants to form virus-like proteins structures. Both candidates were designed for nasal administration and the protein subunit platform was chosen considering its safer profile. For the PanCoV1 and PanCoV2 candidates two main proteins regions, belonging to N and S2 viral antigens, were selected in addition to the RBD. This last antigenic component was included considering its ability as main inducer of neutralizing antibody response; moreover, the Delta strain variant was selected by its higher immunogenicity [15]. By other side, N and S2 proteins are highly conserved among sarbecoviruses, consistently with their crucial role in viral life cycle, and will contribute to a broader vaccine protective scope.

Specifically, PanCoV1 candidate contains the whole N protein as particulate recombinant protein subunit able to induce T cell response mainly. It is known that N protein is essential for encapsulating and packaging the viral RNA genome, forming ribonucleoprotein complexes that are vital for viral assembly and replication. Structurally, the N protein consists of two main domains: the N-terminal domain (NTD), which primarily binds RNA, and the C-terminal domain (CTD), which facilitates dimerization and oligomerization [16,17]. These domains exhibit significant conservation across related coronaviruses, such as SARS-CoV and MERS-CoV, indicating a shared evolutionary lineage and functional importance [18]. The immunogenicity of the N protein has been assessed in various animal models, highlighting its potential as a vaccine candidate [19]. Among the most significant studies involving non-human primates, Hong et al. (2021) evaluated a subunit vaccine in cynomolgus macaques that combined the RBD fused with the tetanus toxoid epitope P2 and the N protein [20]. Their findings revealed that the group immunized intramuscularly with the RBD-P2/N vaccine, along with alum adjuvant, exhibited slightly elevated levels of IgG antibodies, neutralizing antibody titers, and an increased number of IFNγ-secreting cells compared to control formulations. Notably, after challenge with SARS-CoV-2, this bivalent formulation facilitated a faster clearance of the virus compared to the RBD-P2 + alum group. Chiuppesi et al. (2022) demonstrated in AGM that a synthetic multiantigen modified vaccinia Ankara (MVA)-based SARS-CoV-2 vaccine, which co-expresses spike and nucleocapsid antigens (COH04S1), induced robust antigen-specific binding antibodies, neutralizing antibodies, and Th1-biased T cells [5]. This vaccine also provided protection against both upper and lower respiratory tract infections following intranasal/intratracheal challenges with SARS-CoV-2. Similarly, Routhu et al. (2022) reported comparable results in rhesus macaques using an MVA vector expressing both spike and nucleocapsid proteins [21]. They evaluated the immunogenicity and efficacy of this vaccine via intramuscular (IM), buccal, or sublingual routes, finding that the IM vaccination resulted in robust protection after viral challenge, even against heterologous SARS-CoV-2 variants of concern.

In our study, AGM (n=3 per treatment group) were immunized intranasally with three doses of each vaccine preparation. Animals with a pre-existent anti-RBD immunity were part of the study, as a representation of the current worldwide SARS-CoV-2 immunological status, with the majority of the population showing a hybrid anti-SARS-CoV-2 immune response. The different levels of anti-RBD IgG antibodies found in the animals at baseline could reflect the variability presented in the global population.

The PanCoV1 vaccine candidate (N + ODN-39M + RBD) showed a more discrete immunogenicity compared to PanCoV2 (S2NDH + ODN-39M + RBD). However, PanCoV1 preparation successfully induced IgG antibodies against both the RBD and N proteins in two out of three animals after the third dose of vaccine. Additionally, this immune response exhibited cross-reactivity with antigens from SARS-CoV, MERS, and H-CoV. Similar results were observed when assessing mucosal IgA antibodies. Furthermore, PanCoV1 was able to generate a neutralizing antibody response in plasma against SARS-CoV-2 Delta strain virus. This data is consistent with previous results obtained in mice using the VSV-pseudoviral system [8]. Notably, one animal from PanCoV1 treatment group displayed a markedly low immune response during all the study. This animal experienced an increase in respiratory rate after each ketamine administration, however, no other symptoms were observed that would have led to its exclusion from the study. In general, the result obtained after the evaluation of PanCoV1 candidate in AGM are in line with previous data from mice studies [8]. This vaccine candidate induced in BALB/c mice anti-N and RBD cell-mediated immune response in the spleen and also elicited humoral immunity in both sera and lungs. The preparation of PanCoV1 formulation, unlike PanCoV2, includes a centrifugation step following the aggregation reaction with the ODN-39M. This may have contributed to the removal of larger protein aggregates, which could have a higher impact on the immunogenicity of PanCoV1 nasal vaccine preparation in the primate model. This fact, along with the small number of animals employed per treatment group (n=3), could explain the lower immunogenicity observed for PanCoV1 in this study.

On the other hand, the PanCoV2 candidate based on the recombinant protein S2NDH as novel chimeric antigen shows very promising results after its evaluation in AGM. S2NDH protein includes the 800-1020 aa fragment of S2 protein, where a high number of relevant B and T cell epitopes are located [22]. Moreover, it incorporates the C-terminal domain of the nucleocapsid (aa 248-371), which is important for RNA binding and also plays a role in protein dimerization [23]. Several B and T cell epitopes have been also identified within this region, which is highly conserved among sarbecoviruses [22]. PanCoV2 preparation also induced IgG antibodies against RBD, S2 and N proteins in the three intranasally immunized animals after three doses of vaccine. The generated response, shows cross-reactivity with antigens from SARS-CoV, MERS, and H-CoV, and was maintained until the last evaluated time point corresponding with four months after the last dose. The induction of mucosal IgA antibodies was also demonstrated in the three monkeys, mainly against RBD Delta, including a cross-reactive response to RBD and N from SARS-CoV. This was also translated to the generation of higher titres of neutralizing antibody response in plasma against SARS-CoV-2 Delta variant. These results are in accordance with those attained after the evaluation of PanCoV2 intranasal formulation in BALB/c mice. High levels of anti-RBD and anti-N, IgG and IgA antibodies, in both sera and BALF, reactive to proteins from both the SARS-CoV-2 Omicron variant and SARS-CoV, were previously demonstrated [7].

The trend to develop a higher immunogenicity for PanCoV2 candidate could have a potential explanation related with the size of the aggregated structures obtained in the S2NDH + ODN-39M mix. For this specific preparation, after the mixture with the ODN-39M, a more marked pattern of aggregation was found compared with the one observed for PanCoV1 preparations, using transmission electron microscopy (TEM) [7]. This feature intrinsic of PanCoV2 preparation, along with the higher S2NDH+ODN protein dose employed in the present study, could justify the better immune response detected. It is known that the antigen size plays a crucial role in eliciting an effective immune response, particularly following intranasal inoculation. Smaller antigens, typically in the nanoparticle range, are more efficiently taken up by antigen-presenting cells such as dendritic cells (DCs). This efficient uptake is essential for the subsequent activation and maturation of DCs, which are pivotal in initiating T cell responses and facilitating the migration of these cells to lymph nodes [24]. In contrast, larger particles (>500 nm) do not effectively enter lymphatic capillaries without assistance from specialized cells, leading to delayed immune responses [25]. Differences in the antigenic regions included in each vaccine candidate, as well as variations in the aggregation procedure with the ODN-39M, may account for the observed differences in induced immunogenicity in the primate animal model.

The ODN-39M, included as adjuvant in both candidate vaccines, contains CpG motifs designed to activate immune cells in mice, monkeys, and humans [26,27]. Its adjuvant property, along with its capacity to promote aggregation when it is mixed with different viral nucleocapsid proteins, has been previously described [28,29]. Similar to other CpG ODN molecules, it constitutes an agonist of Toll-like receptor 9 (TLR9), a pathogen recognition receptor represented in NALT located dendritic cells [32]. Previous studies from our group have demonstrated that the ODN-39M is able to induce a Th1-biased immune response when is used as adjuvant in protein subunit - based vaccine preparations administered by parenteral or intranasal routes [12,30]. The results obtained in the present study are in line with these previous data.

Another important data coming from this study is the demonstration of the booster capacity of the nasal vaccine candidates under evaluation. They were able to generate high levels of IgG anti-RBD antibodies that persist nearly four months after the last dose. This sustained antibody response is particularly noteworthy, as it suggests that these vaccine candidates could effectively boosts immunity in individuals with pre-existing specific antibodies. Furthermore, the elevated titers of neutralizing antibodies generated in plasma against SARS-CoV-2 Delta strain indicates a robust capacity for viral neutralization, which is critical for protective immunity. These findings underscore the potential of these nasal candidates to elicit a strong and lasting immune response. The generation of a potent neutralizing capacity against SARS-CoV-2 ancestral strain in systemic circulation have been previously demonstrated for both PanCoV candidates in mice studies using a commercial VSV-Spike protein pseudotyped assay. Furthermore, in mice sera both nasal candidates have showed neutralizing capacity againts SARS-CoV [7,8].

On the other hand, considering the classical advantages related with the nasal vaccination and based on the current demonstration of relevant antibody response induced at nasal mucosa after the administration of both PanCoV candidates to AGM, the evaluation of the potential neutralizing capacity in mucosal fluids, as nasal washes, constitutes a very important pending issue. In our previous studies with these two candidates in mice models the neutralizing capacity in BALF samples has been demonstrated [7,8]. Considering this antecedent, a similar behaviour could be expected in AGM model. In the present work the IgA antibody response against RBD was demonstrated in nasal washes and saliva, also with clear evidences of cross-reactive capacity. The demonstration of neutralizing capacity at this compartment could have implications in reduce or cut the viral transmission, which is a crucial goal for respiratory transmitted and highly contagious infections like coronaviruses.

The role of antibody responses and their neutralizing capacity in primary and severe disease with SARS-CoV2 has been proved [31,32]. Although the contribution of T cell response has been more difficult to demonstrate it can also contribute to the immunity with additional mechanisms [33]. In one side, CD4+ T cells provides helper signals to B cells to mount an effective humoral immunity, and additionally, both CD4+ T cells and CD8+ T cells can mediate antiviral functions either by the secretion of cytokines or by the infected cells killing. T cell immunity can be particularly important against the appearance of new variants of concerns in SARS-CoV-2 or new coronaviruses where the cross-reactivity to conserved epitopes could play a more active role or where a larger inoculum is involved and the neutralizing antibody response cannot completely content infection [33]. Even when the response observed in the Placebo group interferes in the analysis, we demonstrated the increase in the number of IFNγ-secreting cells after RBD-Delta stimulation of PBMC from monkeys immunized with both candidates and the maintenance of the response three months after the last dose, with a higher trend to PanCoV2.

The study has some weaknesses that deserve consideration when interpreting its results. A primary concern is the lack of a viral challenge studies, which limits the ability to assess the vaccine’s protective efficacy against CoV infections. The lack of protection studies prevented to test the real contribution of the immune response, including the CMI and mucosal immunity, generated by both PanCoV candidates. So far, the practical issue of BSL3 labs availability to run this experiment has limited our research. Challenge experiments of vaccinated animals using different SARS-CoV-2 VOCs, or even SARS-CoV, to demonstrate the broader protection generate by these nasal candidates should be done in future.

Furthermore, the low number of animals per group may lead to increased variability, limiting the reliability of the conclusions. Taken together, these factors suggest that further research, testing dose levels and different dose administration schedules, is needed to validate the findings and better understand the vaccine’s potential efficacy.

Nevertheless these limitations, we were able to demonstrate that PanCoV nasal candidates, are able to elicit functional humoral and Th1-profiled CMI systemic immunity, mucosal immunity, and cross-reactive humoral immunity in AGM. To our knowledge, this study represents the first evaluation of a Coronavirus broad-scope vaccine candidate based on recombinant proteins administered by intranasal route to non-human primates with previous SARS-CoV-2 immunity.

Specifically, the PanCoV2 formulation (S2NDH + ODN-39M + RBD) demonstrated a remarkable ability to induce cross-reactive immunity against N, S2 and RBD systemically, and also against N and RBD at mucosal compartment, reaching until betacoronavirus level. Notably, it enhanced the immune response to the RBD antigen, leading to improved neutralizing capacity. This PanCoV vaccine emerges as a highly promising candidate for use as a booster alternative, with the potential to amplify and broaden the scope of the immune response generated by previous SARS-CoV-2 exposure, whether through natural infection or vaccination. This innovative approach could play a relevant role in strengthening population immunity against a diverse range of coronaviruses as part of the Pandemic preparedness strategies currently in place worldwide.

## Acknowledgements

The authors extend their special thanks to the researchers at the Civilian Defense Scientific Research Center for their valuable contributions to the microneutralization assays.

## References

[1] Madhav NK, Oppenheim B, Stephenson N, Badker R, Jamison DT, Lam C, et al. Estimated future mortality from pathogens of epidemic and pandemic potential. Center for Global Development; 2023.

[2] Cohen AA, van Doremalen N, Greaney AJ, Andersen H, Sharma A, Starr TN, et al. Mosaic RBD nanoparticles protect against challenge by diverse sarbecoviruses in animal models. Science 2022;377:eabq0839. 10.1126/science.abq0839.

[3] Chang T-Y, Li C-J, Chao T-L, Chang S-Y, Chang S-C. Design of the conserved epitope peptide of SARS-CoV-2 spike protein as the broad-spectrum COVID-19 vaccine. Applied Microbiology and Biotechnology 2024;108:486. 10.1007/s00253-024-13331-y.

[4] Afkhami S, Kang A, Jeyanathan V, Xing Z, Jeyanathan M. Adenoviral-vectored next-generation respiratory mucosal vaccines against COVID-19. Current Opinion in Virology 2023;61:101334. 10.1016/j.coviro.2023.101334.

[5] Chiuppesi F, Nguyen VH, Park Y, Contreras H, Karpinski V, Faircloth K, et al. Synthetic multiantigen MVA vaccine COH04S1 protects against SARS-CoV-2 in Syrian hamsters and non-human primates. NPJ Vaccines 2022;7:7. 10.1038/s41541-022-00436-6.

[6] Leekha A, Saeedi A, Sefat Kmsr, Kumar M, Martinez-Paniagua M, Damian A, et al. Multi-antigen intranasal vaccine protects against challenge with sarbecoviruses and prevents transmission in hamsters. Nat Commun 2024;15:6193. 10.1038/s41467-024-50133-2.

[7] Lobaina Y, Chen R, Suzarte E, Ai P, Musacchio A, Lan Y, et al. A Nasal Vaccine Candidate, Containing Three Antigenic Regions from SARS-CoV-2, to Induce a Broader Response. Vaccines (Basel) 2024;12. 10.3390/vaccines12060588.

[8] Lobaina Y, Chen R, Suzarte E, Ai P, Huerta V, Musacchio A, et al. The Nucleocapsid Protein of SARS-CoV-2, Combined with ODN-39M, Is a Potential Component for an Intranasal Bivalent Vaccine with Broader Functionality. Viruses 2024;16. 10.3390/v16030418.

[9] Hartman AL, Nambulli S, McMillen CM, White AG, Tilston-Lunel NL, Albe JR, et al. SARS-CoV-2 infection of African green monkeys results in mild respiratory disease discernible by PET/CT imaging and shedding of infectious virus from both respiratory and gastrointestinal tracts. PLoS Pathog 2020;16:e1008903. 10.1371/journal.ppat.1008903.

[10] Nambulli S, Escriou N, Rennick LJ, Demers MJ, Tilston-Lunel NL, McElroy AK, et al. A measles-vectored vaccine candidate expressing prefusion-stabilized SARS-CoV-2 spike protein brought to phase I/II clinical trials: protection of African green monkeys from COVID-19 disease. J Virol 2024;98:e0176223. 10.1128/jvi.01762-23.

[11] Woolsey C, Borisevich V, Prasad AN, Agans KN, Deer DJ, Dobias NS, et al. Establishment of an African green monkey model for COVID-19 and protection against re-infection. Nat Immunol 2021;22:86–98. 10.1038/s41590-020-00835-8.

[12] Gil L, Marcos E, Izquierdo A, Lazo L, Valdés I, Ambala P, et al. The protein DIIIC-2, aggregated with a specific oligodeoxynucleotide and adjuvanted in alum, protects mice and monkeys against DENV-2. Immunol Cell Biol 2015;93:57–66. 10.1038/icb.2014.63.

[13] Gil L, Lazo L, Valdés I, Suzarte E, Yen P, Ramírez R, et al. The tetravalent formulation of domain III-capsid proteins recalls memory B- and T-cell responses induced in monkeys by an experimental dengue virus infection. Clin Transl Immunology 2017;6:e148. 10.1038/cti.2017.24.

[14] Manenti A, Maggetti M, Casa E, Martinuzzi D, Torelli A, Trombetta CM, et al. Evaluation of SARS-CoV-2 neutralizing antibodies using a CPE-based colorimetric live virus micro-neutralization assay in human serum samples. J Med Virol 2020;92:2096–104. 10.1002/jmv.25986.

[15] Aparicio B, Lasarte JJ, Sarobe P. Mutated trimeric RBD vaccines: a platform against variants of concern. Signal Transduct Target Ther 2023;8:161. 10.1038/s41392-023-01426-3.

[16] Chen C-Y, Chang C-K, Chang Y-W, Sue S-C, Bai H-I, Riang L, et al. Structure of the SARS coronavirus nucleocapsid protein RNA-binding dimerization domain suggests a mechanism for helical packaging of viral RNA. J Mol Biol 2007;368:1075–86. 10.1016/j.jmb.2007.02.069.

[17] Cubuk J, Alston JJ, Incicco JJ, Singh S, Stuchell-Brereton MD, Ward MD, et al. The SARS-CoV-2 nucleocapsid protein is dynamic, disordered, and phase separates with RNA. Nat Commun 2021;12:1936. 10.1038/s41467-021-21953-3.

[18] Huang Y, Chen J, Chen S, Huang C, Li B, Li J, et al. Molecular characterization of SARS-CoV-2 nucleocapsid protein. Front Cell Infect Microbiol 2024;14:1415885. 10.3389/fcimb.2024.1415885.

[19] Song W, Fang Z, Ma F, Li J, Huang Z, Zhang Y, et al. The role of SARS-CoV-2 N protein in diagnosis and vaccination in the context of emerging variants: present status and prospects. Front Microbiol 2023;14:1217567. 10.3389/fmicb.2023.1217567.

[20] Hong S-H, Oh H, Park YW, Kwak HW, Oh EY, Park H-J, et al. Immunization with RBD-P2 and N protects against SARS-CoV-2 in nonhuman primates. Sci Adv 2021;7. 10.1126/sciadv.abg7156.

[21] Routhu NK, Gangadhara S, Lai L, Davis-Gardner ME, Floyd K, Shiferaw A, et al. A modified vaccinia Ankara vaccine expressing spike and nucleocapsid protects rhesus macaques against SARS-CoV-2 Delta infection. Sci Immunol 2022;7:eabo0226. 10.1126/sciimmunol.abo0226.

[22] Agarwal A, Beck KL, Capponi S, Kunitomi M, Nayar G, Seabolt E, et al. Predicting Epitope Candidates for SARS-CoV-2. Viruses 2022;14. 10.3390/v14081837.

[23] Chang C, Hou M-H, Chang C-F, Hsiao C-D, Huang T. The SARS coronavirus nucleocapsid protein--forms and functions. Antiviral Res 2014;103:39–50. 10.1016/j.antiviral.2013.12.009.

[24] Kehagia E, Papakyriakopoulou P, Valsami G. Advances in intranasal vaccine delivery: A promising non-invasive route of immunization. Vaccine 2023;41:3589–603. 10.1016/j.vaccine.2023.05.011.

[25] Bachmann MF, Jennings GT. Vaccine delivery: a matter of size, geometry, kinetics and molecular patterns. Nat Rev Immunol 2010;10:787–96. 10.1038/nri2868.

[26] Verthelyi D, Ishii KJ, Gursel M, Takeshita F, Klinman DM. Human peripheral blood cells differentially recognize and respond to two distinct CPG motifs. J Immunol 2001;166:2372–7. 10.4049/jimmunol.166.4.2372.

[27] Krug A, Rothenfusser S, Hornung V, Jahrsdörfer B, Blackwell S, Ballas ZK, et al. Identification of CpG oligonucleotide sequences with high induction of IFN-alpha/beta in plasmacytoid dendritic cells. Eur J Immunol 2001;31:2154–63. 10.1002/1521-4141(200107)31:7<2154::aid-immu2154>3.0.co;2-u.

[28] Valdes I, Gil L, Lazo L, Cobas K, Romero Y, Bruno A, et al. Recombinant protein based on domain III and capsid regions of zika virus induces humoral and cellular immune response in immunocompetent BALB/c mice. Vaccine 2023;41:5892–900. 10.1016/j.vaccine.2023.08.035.

[29] Gil L, Izquierdo A, Lazo L, Valdés I, Ambala P, Ochola L, et al. Capsid protein: evidences about the partial protective role of neutralizing antibody-independent immunity against dengue in monkeys. Virology 2014;456–457:70–6. 10.1016/j.virol.2014.03.011.

[30] Lazo Vázquez L, Gil González L, Marcos López E, Pérez Fuentes Y, Cervetto de Armas L, Brown Richards E, et al. Evaluation in Mice of the Immunogenicity of a Tetravalent Subunit Vaccine Candidate Against Dengue Virus Using Mucosal and Parenteral Immunization Routes. Viral Immunol 2017;30:350– 8. 10.1089/vim.2016.0150.

[31] O’Brien MP, Forleo-Neto E, Musser BJ, Isa F, Chan K-C, Sarkar N, et al. Subcutaneous REGEN-COV Antibody Combination to Prevent Covid-19. N Engl J Med 2021;385:1184–95. 10.1056/NEJMoa2109682.

[32] Khoury DS, Cromer D, Reynaldi A, Schlub TE, Wheatley AK, Juno JA, et al. Neutralizing antibody levels are highly predictive of immune protection from symptomatic SARS-CoV-2 infection. Nat Med 2021;27:1205–11. 10.1038/s41591-021-01377-8.

[33] Kent SJ, Khoury DS, Reynaldi A, Juno JA, Wheatley AK, Stadler E, et al. Disentangling the relative importance of T cell responses in COVID-19: leading actors or supporting cast? Nat Rev Immunol 2022;22:387–97. 10.1038/s41577-022-00716-1.

